# Within-ejaculate sperm selection in humans acts against molecular signatures of age-related disease

**DOI:** 10.1101/2024.10.08.617222

**Authors:** Daniel Marcu, Jayme Cohen-Krais, Alice Godden, Ghazal Alavioon, Anders Bergström, Kristian Almstrup, Alexei A. Maklakov, Gerhard Saalbach, Simone Immler

## Abstract

The germline is widely regarded as a checkpoint against the inheritance of damaged genomes, yet the mechanisms that could enact such filtering remain poorly resolved. Of the millions of sperm in a human ejaculate, only one fertilises the egg, creating a strong opportunity for selection among gametes. Combining within-ejaculate selection on sperm quality with whole-genome sequencing and proteomics in healthy donors, we find that this selection is biased against molecular signatures of age-related disease. The most reproducibly diverging genes are tumour suppressors, and genes diverging under sperm longevity-based selection are enriched for senescence-associated genes involved in genome maintenance and oxidative stress response. High-quality sperm are further depleted of inflammation- and cancer-associated proteins. This genomic signature is conserved in zebrafish, in which longer-lived sperm sire longer-lived offspring. We propose that within-ejaculate selection acts as a pre-fertilisation filter against age-related disease alleles, with implications for offspring lifespan and healthspan.

## Introduction

How the germline avoids transmitting damaged genomes, including those that predispose to age-related disease in the next generation, is not well understood. One underexplored possibility is pre-fertilisation selection among gametes. A human ejaculate contains between 20 and 150 million sperm, of which one fertilises an egg. Sperm within the ejaculate of a man vary both phenotypically and genetically, offering a strong opportunity for natural selection to act upon (1–3). Although the importance of selection among germ cells is increasingly recognised (4), the role and magnitude of selection among mature sperm within an ejaculate, acting after spermatogenesis is complete and before fertilisation, remains largely overlooked (1,3). Recent findings in zebrafish (*Danio rerio*) show that longer-lived sperm differ from shorter-lived sperm at many loci distributed across the genome (5). Large-scale studies of gene expression during spermatogenesis provide clear evidence for expression in post-meiotic haploid spermatids (6), and the incomplete sharing of transcripts between meiotic products (7) supports the idea that genetic differences among sperm engender phenotypic differences within an ejaculate. This notion is further supported by accelerated evolution of genes expressed at post-meiotic stages (6), as well as elevated purifying selection on these genes (8).

Despite this, the genes and associated molecular markers under haploid selection in sperm are not known. Identifying molecular biomarkers of high-quality sperm is central to understanding the fundamental biology of human reproduction and fertility and will also inform improved sperm selection for use in Assisted Reproductive Technology (9,10). If selection among sperm acts on the genetic and proteomic state of the gamete, the alleles and proteins that survive it should be biased away from those that compromise long-term cellular function and toward those that sustain it - a prediction that, to our knowledge, has not been tested at the molecular level in any vertebrate.

Here, we designed two in vitro assays (swim up and methyl cellulose; see Materials and Methods) to select for high-quality sperm within the ejaculates of human donors and combined this selection with deep-coverage whole-genome sequencing (WGS) and tandem mass tag (TMT) proteomics to resolve the molecular architecture of human sperm quality, and to ask whether pre-fertilisation selection acts against the genetic and molecular signatures of age-related disease. Complementary analyses using polygenic scores from the PAN-UK Biobank further enabled assessment of putative implications of selection on sperm quality variants for metabolic ageing and late-life disease, such as cancer. By comparing the genomic signature of haploid selection in human sperm with that in zebrafish, in which longer-lived sperm sire longer-lived offspring (41), we assessed the biological relevance and generality of our findings across taxa.

## Results

### Selection on phenotypic sperm quality

We selected distinct sperm-quality pools from semen samples of 35 donors using the swim-up assay and compared fertilisation-relevant traits (sperm kinematics, morphology, chromatin, DNA integrity) across raw and washed samples and selected sperm incubated for 0, 4 and 24 hours (Suppl. Table S1). The fraction of motile sperm decreased from an average of 60% in fresh and washed samples to 50% after 4 hours of incubation and 25% after 24 hours of incubation prior to swim-up (Suppl. Fig. S1 and Suppl. Table S2). Sperm kinematics improved continuously with increasing incubation time (Fig. 1A; Suppl. Tables S3-S10; Suppl. Fig. S2), as did chromatin condensation (Fig. 1B; Suppl. Table S11) and gross morphology (Fig. 1C; Suppl. Table S12), whereas DNA integrity was slightly higher in swim-up samples compared to all other samples (Fig. 1D; Suppl. Table S13).

**Figure 1:**
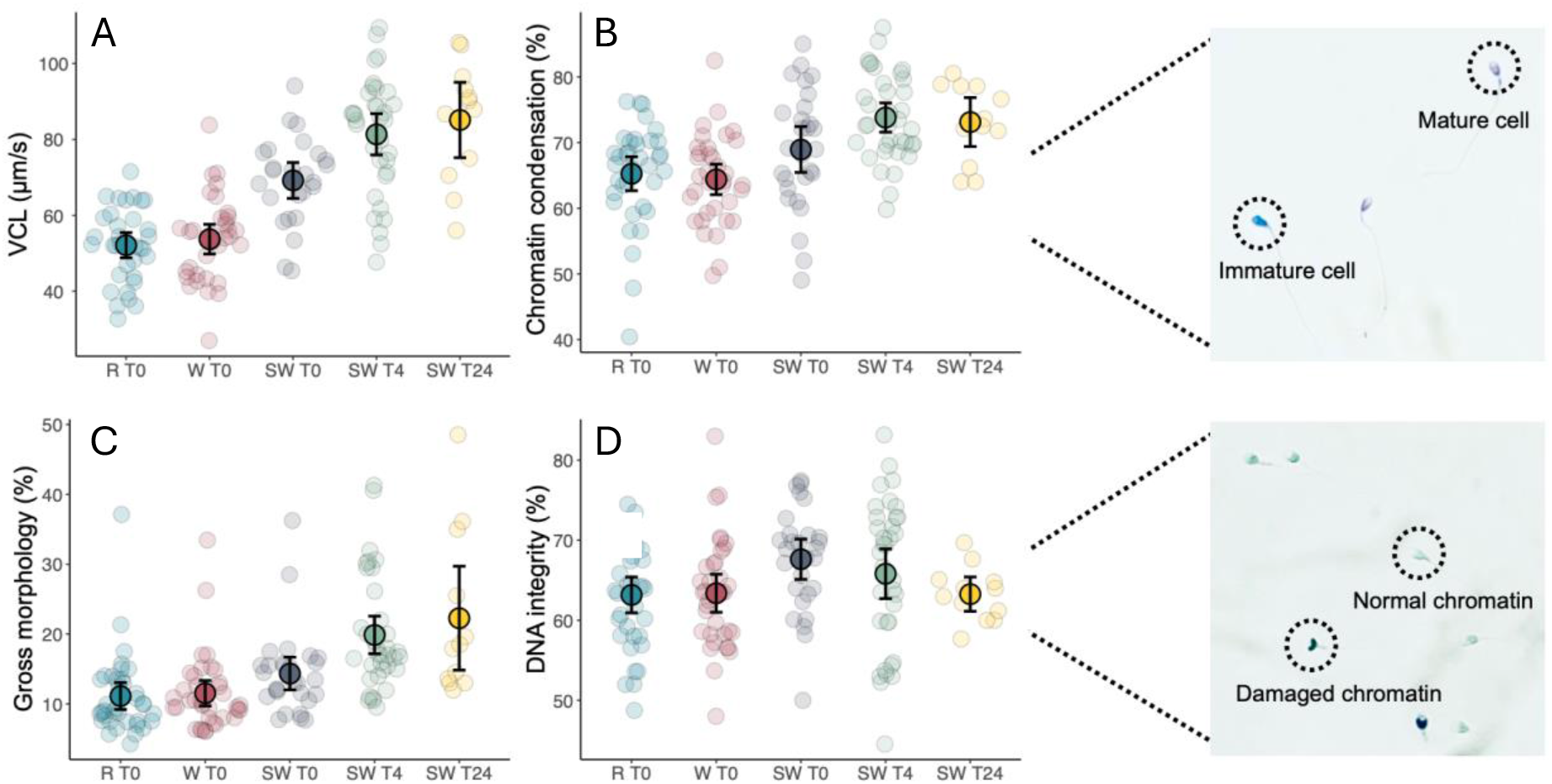
Phenotypic signature of within-ejaculate selection on sperm quality. Phenotypic traits of sperm in raw (R T0), washed (W T0), swim up after 0 hours incubation (SW T0), after four hours incubation (SW T4) and 24 hours incubation (SW T24). (A) Curvilinear velocity, (B) % sperm with mature chromatin (using aniline), (C) % sperm with normal morphology (using SpermBlue), and (D) % sperm without single- or double-strand breaks (using toluidine blue). Data for all donors are shown as individual dots and overall mean ± standard error are shown in black for each time point (sample sizes: Raw: n=35; W: n=35; T0: n=25; T4: n=35 and T24: n=12).

### Genomic divergence among selected sperm pools

We linked phenotypic differences between selected high- and low-quality sperm with genomic differences by performing high-coverage WGS on selected sperm pools using both assays. For the methyl cellulose assays, we used ejaculates from four donors and collected sperm from the Centre (low-quality) and the Edge (high-quality) of the incubation dish; for the swim-up assays, we used ejaculates from five donors and collected sperm at 0 hours (T0; low-quality) and at 4 and 24 hours (T4 and T24; high-quality). Each donor represented an experimental replicate and samples were analysed separately. For DNA extraction and sequencing, we standardised the number of cells to 2 million per sample and performed WGS at ∼100× per sample.

To assess genetic variation across donors, we conducted a comprehensive analysis of allele frequencies and identified significant loci for interpretation. We compared allele frequencies at an average of 2,280,804 loci per donor (min = 2,022,290; max = 2,754,360; Suppl. Table S14) and retained >99.9th-quantile significant loci for interpretation. We identified heterozygous sites for each donor, filtered for a minimum total read number of 32 per locus, and performed likelihood-ratio tests on allele frequencies to test for significant deviations from expected Mendelian frequencies using the Centre/T0 samples as reference. We identified significant variants at an average of 1,139 loci per donor (min = 783; max = 2,578; Fig. 2A,B; Suppl. Table S15 & S16; Suppl. Fig. S10; coverage was not correlated with significance; Suppl. Table S17; Suppl. Fig. S3)).

**Figure 2:**
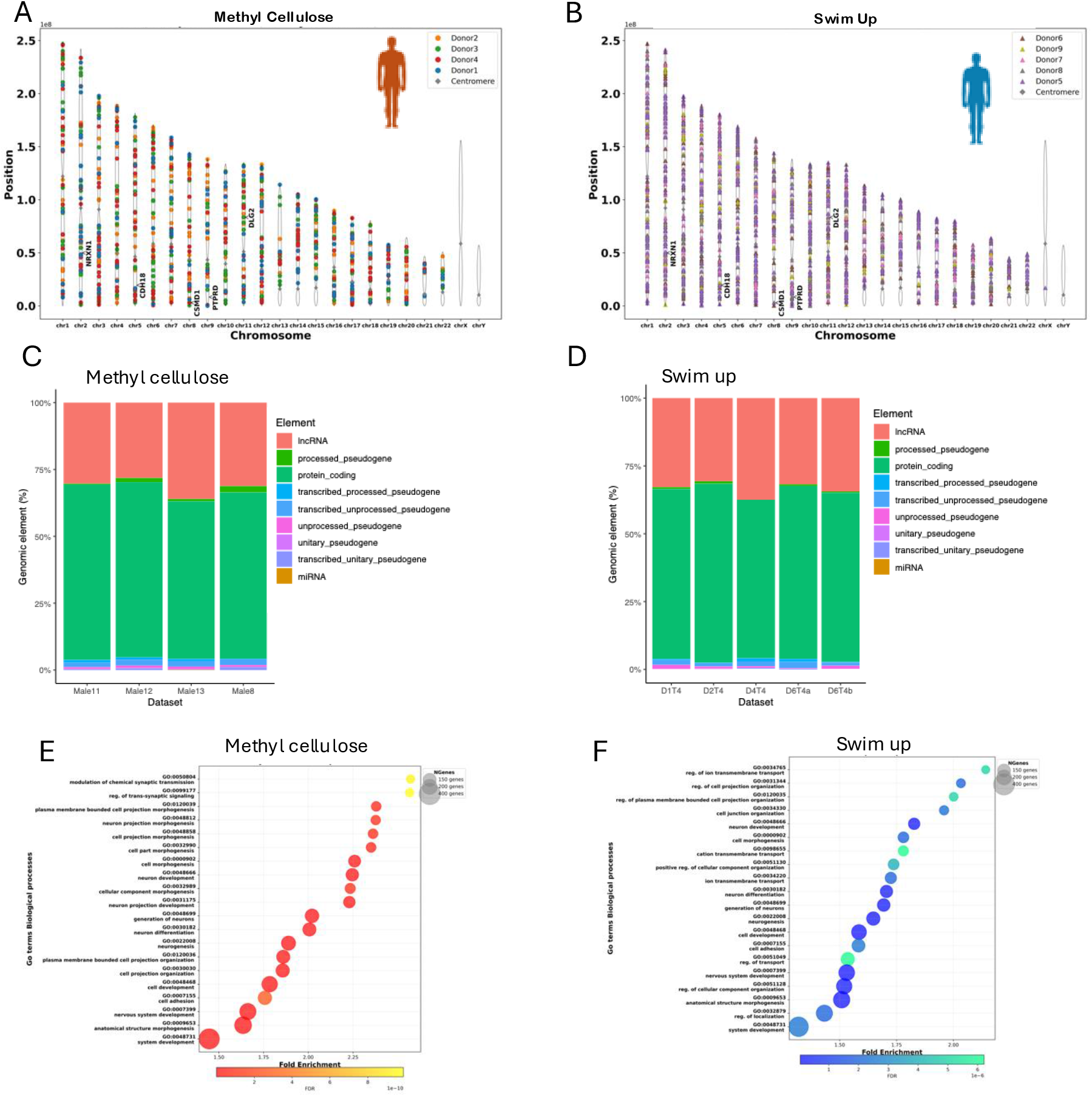
Genomic signature of within-ejaculate selection on human sperm quality. Loci with significantly diverging alleles between sperm pools collected (A) from Centre (low-quality) and Edge (high-quality) in the Methyl Cellulose assay (n = 4) and (B) at T0 (low quality) and T4 (high quality) from Swim Up assays (n = 5; likelihood ratio tests, > 99.9th quantile). Top 20 significant Biological GO terms for the genes showing significant divergence in (C) Methyl cellulose and (D) Swim Up. Distribution of genomic elements among significantly diverging loci for (E) Methyl Cellulose and (F) Swim Up.

We mapped the significantly diverging loci to an average of 511 genetic elements (min = 340; max = 697) per comparison, of which 64% were intron and exon regions of protein-coding genes (min = 58.4%; max = 69.4%) and 31.6% were lncRNAs (min = 26.4%; max = 37.5%; Fig. 2C,D), with no significant differences in element representation across donors and time points (χ² = 103.42, df = 228, p = 1.0). Significantly diverging protein-coding sequences were longer than the average protein-coding gene in the reference genome (Centre: t = 8.62, p < 0.001; T4: t = 5.66, p < 0.001; T24: t = 7.42, p < 0.001; T48: t = 4.42, p < 0.001; Suppl. Fig. S4). Gene functions and Gene Ontology (GO) terms overlapped across time points (Suppl. Fig. S5-S7); we focus here on methyl cellulose and swim up at T4. Overall, significantly diverging genes were enriched for processes of cell assembly and neuronal signalling. In methyl cellulose, enrichment included transmembrane receptor protein tyrosine kinase signalling, cell junction assembly and neuron development (Fig. 2E), and molecular functions associated with cell communication and membrane transport such as protein serine kinase activity, G-protein-coupled neurotransmitter receptor activity, calcium ion binding and glutamate receptor activity (Suppl. Fig. S6). In swim up at T4, most enrichment was associated with nervous system development, including synapse assembly, neuron differentiation and chemical synaptic transmission (Fig. 2F), and molecular processes involving cellular transport and cell adhesion such as calcium ion binding, glutamate receptor activity and calcium-activated potassium channel activity (Suppl. Fig. S7).

Across both assays and all time points, we identified 19 genes that diverged in at least six of nine donors (Suppl. Table S18); the top five were *DLG2*, *PTPRD*, *CSMD1*, *CDH18* and *NRXN1*. *DLG2*, *PTPRD*, *CSMD1* and *CDH18* are well-described tumour suppressor genes (11–13), and NRXN1 has a role in lung cancer development. All five genes have also been associated with altered risk for schizophrenia, autism spectrum disorder (ASD), and neurodevelopmental disorders, as well as involvement in inflammasome formation and cellular senescence (14–18). *CSMD1* is associated with male and female infertility (19), *DLG2* is downregulated in human sperm with compromised performance (20), CDH18 is implicated in cell adhesion in testes (21) and deletions in *NRXN1* have been observed in azoospermic patients (22). Analysing assays separately, we detected *RBFOX1*, *EGLN3*, *CDH13*, *LRP1B* and *LINC00960* across all four donors in methyl cellulose (Suppl. Fig. S8A) and *CDH18* and *ZFPM2* across all donors in swim up at T4 (Suppl. Fig. S8B). *RBFOX1* and *CDH13* contribute to neuronal development (23,24), whereas *CDH18*, *EGLN3*, *LRP1B* and *LINC00960* participate in cancer cell migration and tumour growth or suppression (25–27). None of these variants are currently associated with known phenotypes. We therefore compared all significantly diverging variants in our data against the UK BioBank dataset and identified four variants with known associated phenotypes in Methyl Cellulose data and 12 for Swim Up assays. We compared the minor allele frequencies of these known variants in unselected and selected pools and observed a trend of reduced frequencies for most variants in the selected pool (two out of four in MC and 11 out of 12 in SU; Fig. 3A,B). In addition, we tested direction of selection for variants in CellAge associated with stress-response genes predominantly: they exhibited negative allele-frequency shifts (ΔMaf = Maf_O − Maf_C), indicating depletion of alternative alleles in the selected sperm pool. Among the five variants with available CADD annotation, four showed decreases in allele frequency, whereas a single variant increased in frequency (Fig. 3E). Notably, the only variant classified as deleterious by CADD PHRED score (CADD PHRED = 21.4; top ∼1% of predicted deleterious variants genome-wide) also demonstrated a substantial reduction in frequency (ΔMaf ≈ −0.35), consistent with purifying selection against a potentially functional allele. In contrast, variants with lower predicted deleteriousness (CADD PHRED <10) showed both positive and negative frequency shifts. Overall, selecting for higher quality sperm has the potential of reducing the frequency of deleterious alleles in the sperm pool.

**Figure 3:**
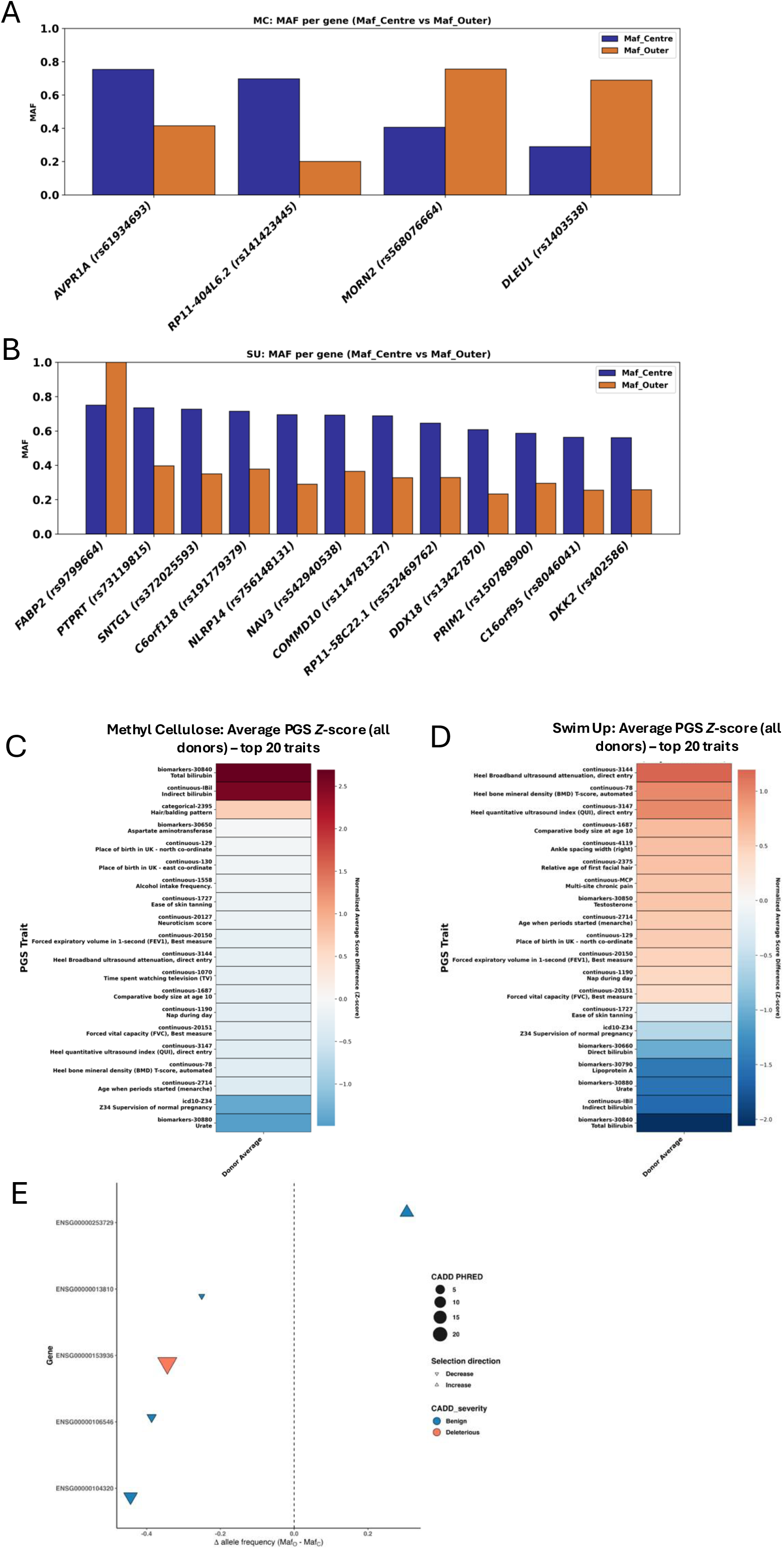
Biological functions and direction of selection in variants diverging between high- and low-quality human sperm pools. Significantly diverging variants based on likelihood ratio tests (padj < 0.00001) listed as known variants with associated phenotypes on UK Biobank with direction of selection indicated by Minor Allele Frequency (MAF) between unselected (blue) and selected (red) sperm pools for (A) Methyl Cellulose and (B) Swim Up assays. Heatmap showing Z-score normalised average differences in polygenic scores for the top traits selected based on the 99th percentile of absolute score differences across donors. Positive values (red) indicate traits with higher-than-average score differences, while negative values (blue) indicate lower-than-average differences for (G) Methyl Cellulose and (H) Swim Up.

### Polygenic scores of variants diverging between sperm pools

We connected the genomic signature of selection for sperm quality with traits by generating polygenic scores (see Material & Methods) using the Pan-UK Biobank database (43). Using raw PGS that included all heterozygous loci for each sperm pool (T0/T4 and Centre/Edge), we calculated the score difference between the two pools for each donor and normalised the difference. Based on normalised Z-scores in the 99th percentile, we identified the highest-scoring traits as total, direct and indirect bilirubin, supervision of normal pregnancy, lipoprotein(a) and urate and some lifestyle and diet traits were also among the top traits (Fig. 3C,D). The emerging terms were remarkably consistent across donors (Suppl. Fig. S9), and changes in polygenetic scores for bilirubin-related traits showed a negative correlation to changes in scores for lifestyle traits, indicating genetic correlation between traits. The direction of effects was not always consistent between donors and experiments. This supports the idea that selection on sperm performance is likely polygenic, rather than driven by a small number of variants. In addition, the variability across genomic backgrounds may indicate potential epistatic interactions, although polygenicity alone does not always imply epistasis. We tested for genetic correlations between trait-associated polygenic scores, which yielded similar patterns across both assays. (Suppl. Fig. S10). Overall, these results support a polygenic response to sperm selection, and that organism-level traits have genetic underpinnings that are visible to selection at the sperm stage.

### Evolutionarily conserved genomic signature of haploid selection between humans and zebrafish

We re-analysed zebrafish sperm-pool data from Alavioon et al. (2017) (5) using the same rigorous approach as for the human data and compared genes diverging between sperm pools in humans and zebrafish using human gene annotations (Suppl. Tables S19-S21; Suppl. Fig. S11-S12). We identified on average 5,233 significantly diverging loci per male (range: 4,545-5,722; Fig. 4A), with distributions of genomic elements like those observed in human samples (Fig. 4B). Enrichment analyses in zebrafish revealed biological processes including cell-to-cell interactions and neurodevelopmental processes (differentiation and generation of neurons, cell signalling, developmental cell differentiation and organ formation; Fig. 4C), and molecular functions related to ion transport and channel activity (calcium-activated cation channel activity, sodium channel activity and ion transmembrane transporter activity; Suppl. Fig. S13).

**Figure 4:**
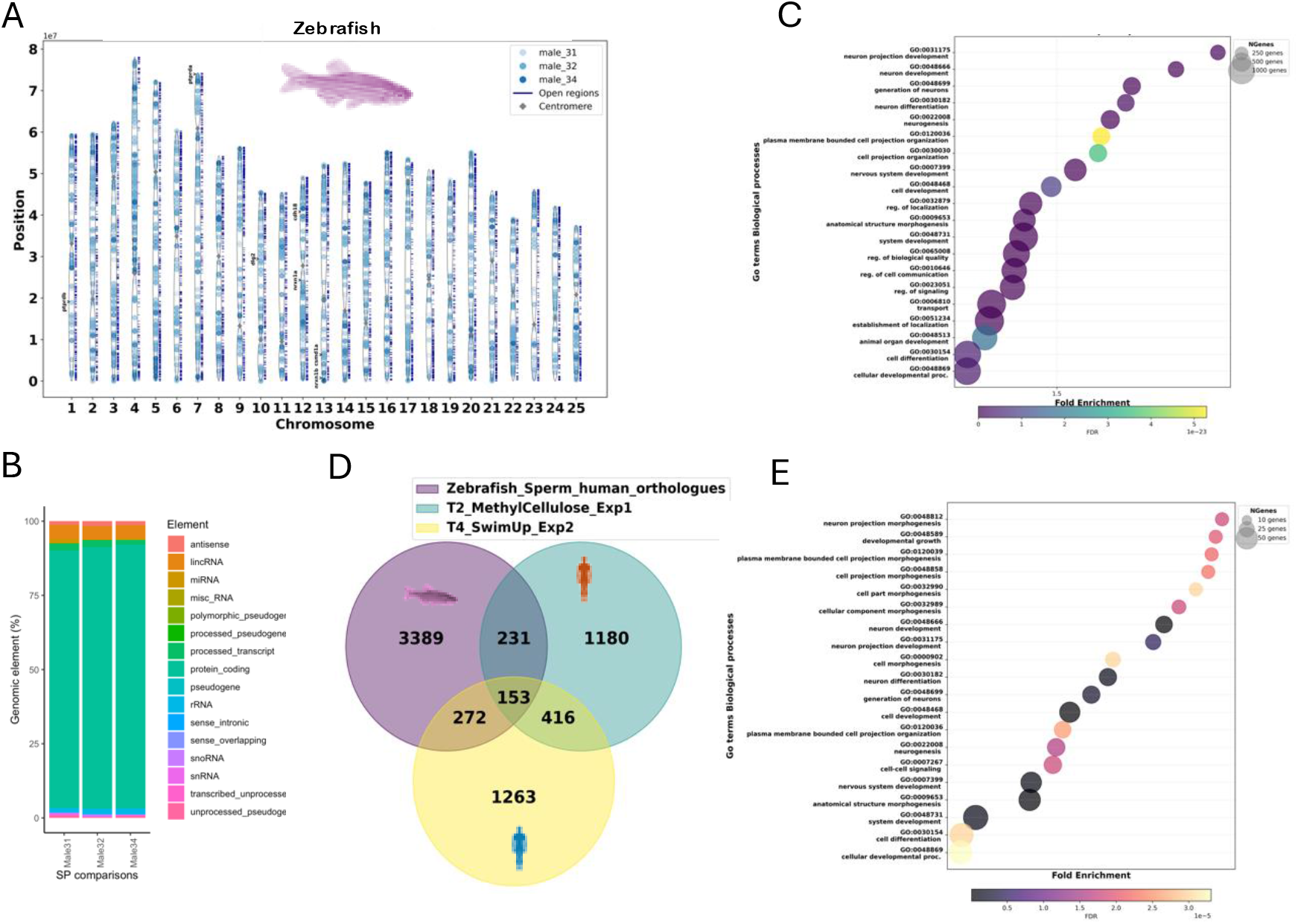
Comparison of genomic signature of within-ejaculate selection in zebrafish and humans. (A) Loci with significantly diverging alleles between sperm pools collected from centre (low quality) and edge of a water droplet (n = 3; data from Alavioon et al 20175) and (B) the associated top 20 significant biological GO terms. (C) Distribution of genomic elements among significantly diverging loci. (D) Venn diagram showing overlap of significantly diverging genes between human Methyl Cellulose, human Swim Up and zebrafish. (E) Top 20 significant biological GO terms for the 153 genes share between zebrafish, human Methyl Cellulose and human Swim Up assays.

We found 153 genes shared among human methyl cellulose, swim up and the human homologs of zebrafish genes (Fig. 4D). The top 20 significant biological GO terms for these 153 genes indicated functions centred on neural development and cell proliferation (Fig. 4E). Semantic similarity analysis between GO terms identified in humans and the human homologs of zebrafish genes revealed high similarity (scores of 0.85 for methyl cellulose and 0.7 for swim up at T4; Suppl. Fig. S14). Finally, three of the five top candidate genes in humans were also significantly diverging among zebrafish sperm pools - *DLG2*/*dlg2*, *CSMD1*/*csmd1a* and *CDH18*/*cdh18*. Altogether, despite ∼450 million years of evolutionary divergence between zebrafish and humans (28), the biological processes and gene functions under haploid selection in sperm appear evolutionarily conserved across taxa.

### Proteomic biomarkers of selected sperm

To identify molecular biomarkers of sperm quality, we performed methyl cellulose and swim up assays in an additional four human donor samples (two donors per assay) and applied TMT proteomics to the selected sperm samples (see Material & Methods), including a technical replicate for one donor to confirm robustness. Of 8,191 peptide matches, 4,616 proteins were identified and retained after rigorous filtering, and 2,232 overlapped across all donors (Fig. 5A). In total, 1,178 proteins showed significant differences in abundance between sperm pools in at least one donor, with 634 proteins showing significantly (log2-fold change > 2; *p_adj_* < 0.05) lower abundance and 544 showing significantly higher abundance in high-quality sperm (Fig. 5B). Analysing assays separately, we identified 52 and 425 proteins with higher and lower abundance, respectively, in methyl cellulose and 33 and 594 proteins with higher and lower abundance, respectively, in swim up (range: 43-113 higher and 86-431 lower per donor; Suppl. Table S22). In swim up, proteins less abundant in high-quality sperm were associated with RNA processing and protein translation (Suppl. Fig. S15A), whereas proteins more abundant in high-quality sperm were enriched for motility and membrane organisation. In methyl cellulose, proteins were associated with lipid transport and cholesterol efflux pathways characteristic of capacitation (Suppl. Fig. S15B). KEGG analysis showed that proteins less abundant in selected sperm were enriched for RNA- and ribosome-binding activities, including “ribonucleoprotein complex binding” and “structural constituent of ribosome” (Suppl. Fig. S16A). Adhesion-related terms such as “collagen binding” and “cell-substrate junction” were also observed. Proteins more abundant in selected sperm were enriched for functions related to enzyme inhibition and structural support, including endopeptidase inhibitor activity, peptidase regulator activity and antioxidant activity (Suppl. Fig. S16B). Functional enrichment analyses revealed two broad, consistent patterns. First, proteins with low abundance in high-quality sperm are linked to translational and splicing activity - features of immature sperm absent from mature, high-quality sperm (29). Their depletion in selected fractions supports the interpretation that sperm surviving the selection assays represent a more functionally mature population. Second, proteins with increased abundance in high-quality sperm were related to motility, consistent with evidence that flagellar structure and activity are central to sperm motility and fertilisation competence (30–32). Other enriched functions included lipid metabolism and membrane-associated protein complexes, processes tightly linked to capacitation and regulation of membrane fluidity (33). Enrichment for antioxidant and enzyme inhibitor activities is also consistent with requirements to withstand oxidative and proteolytic stress during transit through the female tract (34,35). Applying an additional filtering step based on significance (*p_adj_* < 0.05) and the median ratio of protein abundance among sperm pools across all four donors, we identified six key proteins that were downregulated and located on the membrane in high-quality sperm: OLFM4, CTNG, PLA2, DEFB105A, midkine and protein K, all associated with human disease phenotypes including various cancers, inflammation and cell growth 36-40. We validated OLFM4 and CTNG as possible biomarkers based on their abundance and putative cell-surface localisation using fluorophore-conjugated antibodies and FACS in five additional donors (see Methods). Both proteins showed consistently higher abundance in low-quality sperm across all five donors (Fig. 5C; Suppl. Table S23; Suppl. Fig. S17, S18), supporting the TMT results. Genes underlying the significantly diverging proteins showed some overlap with genomic signatures in the WGS data.

**Figure 5:**
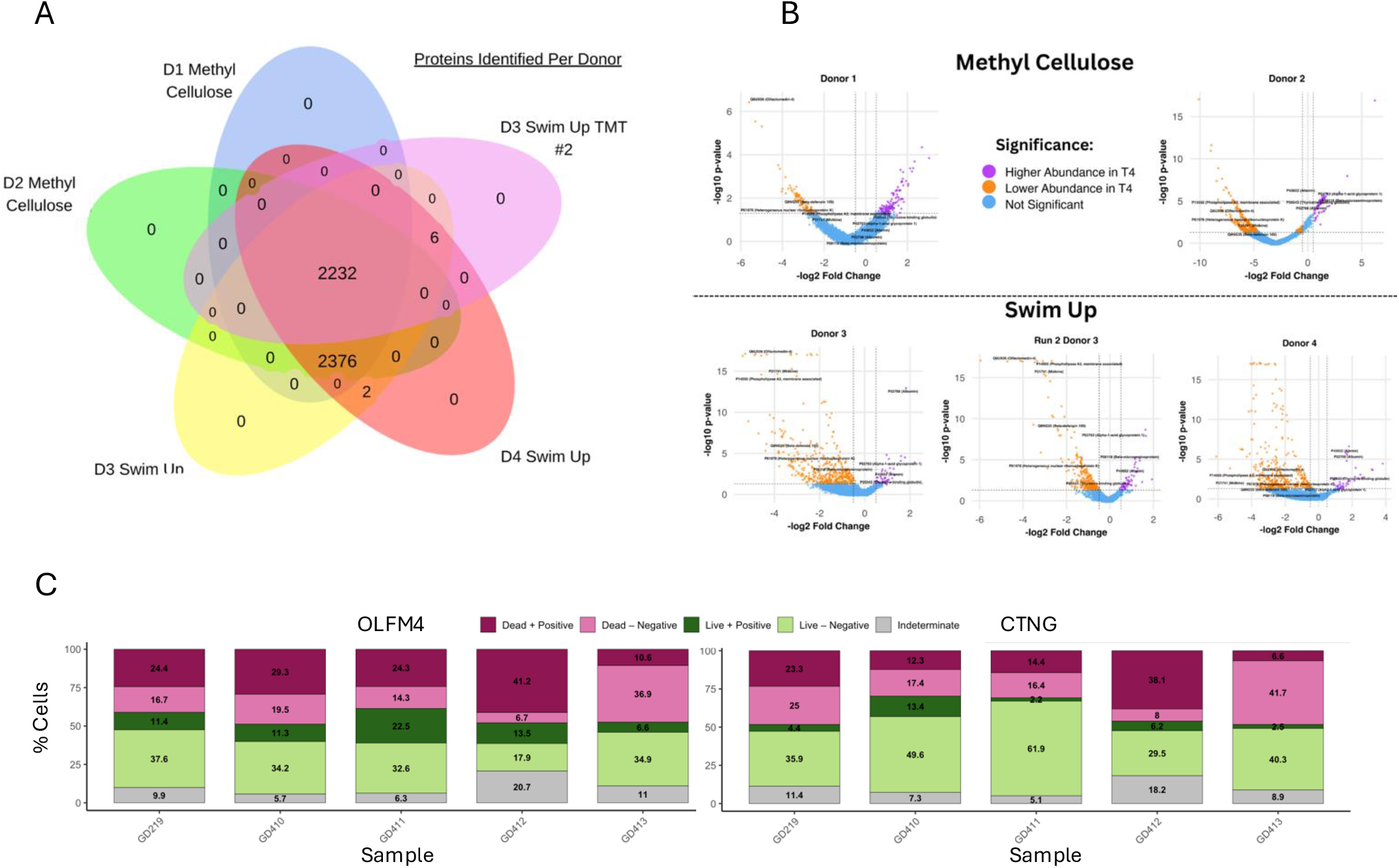
Proteomic signature of within-ejaculate selection on sperm performance. (A) Venn diagram showing overlap of differentially expressed proteins between donors, experiments and technical runs. (B) Volcano plots for each donor displaying significantly differentially expressed proteins (pink; (significance cut-off points are padj < 0.05; log2 fold change > |2|) between Centre/T0 and Edge/T4 sperm pools assays with log2 fold changes > 0 indicating increased expression in Centre/T0 sperm pools and log2 fold changes < 0 indicating increased expression in Edge/T0 sperm pools. Two technical runs were performed for Donor A in the Swim Up assay. The top five significant proteins shared among all samples are highlighted by black circle and protein ID. (C) Validation of OLFM and CTNG as markers of human sperm quality using FACS on cells stained with a combination of antibody labelling and live-dead staining. ‘Positive’ indicates staining for marker protein above gating threshold and ‘negative’ indicates sub-threshold labelling for antibodies (see Supplementary Material for gating details).

## Discussion

The data presented here support a model in which within-ejaculate sperm selection acts as a pre-fertilization filter biased against the molecular hallmarks of age-related disease. Three lines of evidence converge on this interpretation. First, the genes most reproducibly diverging between high- and low-quality sperm - *DLG2* (11), *PTPRD* (12,13), *CSMD1* (19), *CDH18* (21) and *NRXN1* (14–18), supplemented by *RBFOX1* (23), *EGLN3* (25), *CDH13* (24), *LRP1B* (26) and *LINC00960* (27) in single-assay analyses - are dominated by tumour suppressors and modulators of cellular senescence and inflammasome formation. Cancer is the prototypical age-related disease, and loss of tumour-suppressor function is a classical early step in oncogenesis. Biased depletion of variants in these genes from successful sperm pools is therefore consistent with a quality-control function operating against the genetic basis of late-life disease. Second, polygenic score divergence between sperm pools concentrated on canonical metabolic ageing biomarkers, such as lipoprotein(a) and urate, alongside scores related to lifestyle and diet. Lipoprotein(a) and urate are robust predictors of cardiovascular and metabolic mortality (55,56), and their emergence here suggests that the polygenic architecture of human metabolic ageing is partly visible to selection at the gametic stage. Third, proteomic profiling identified 1,178 differentially abundant proteins between high- and low-quality sperm. In high-quality sperm, cell-surface proteins implicated in inflammation, immune dysregulation and tumour progression, including OLFM4, midkine and a secreted phospholipase A2 (36,38,39), were significantly depleted, whereas proteins associated with motility, antioxidant defence and proteolytic resilience were enriched. Together, these patterns indicate that the gamete that fertilizes the oocyte is a non-random selection biased toward genomes carrying fewer alleles, and proteomes displaying fewer states, associated with age-related disease.

A complementary line of evidence focuses on the role of within-ejaculate selection in relation to a specific molecular pathway. Genes diverging under longevity-based (swim up) selection were enriched for curated regulators of cellular senescence (CellAge, 57), an enrichment absent under motility based (methyl cellulose) selection. Within this gene subset, the recovered genes were predominantly those whose primary function is genome maintenance and the detoxification of oxidative damage, specifically the DNA double-strand break sensors NBN and PRKDC (58, 59) and the reactive aldehyde and redox enzymes ALDH2 and ME1 (60, 61). These genes are catalogued as senescence regulators because their loss precipitates senescence in somatic cells, but their underlying role is to preserve genome integrity and buffer oxidative stress. This class of genes is mechanistically very relevant to the selection imposed, since a swim up treatment subjects sperm to sustained oxidative and metabolic stress, and sperm carrying compromised genome maintenance or antioxidant machinery would be expected to fail such a challenge. At these genes, alleles were consistently depleted in high-quality sperm, indicating directional selection. However, the diverging variants were overwhelmingly non-coding and carried no signature of predicted protein-damaging consequence, pointing to selection acting on the regulation of these genes rather than on coding deleterious alleles. We therefore propose that within-ejaculate selection favours better genome maintenance and stress resistance, offering a candidate mechanism linking gametic selection to organismal lifespan and healthspan.

The genomic markers of haploid selection are remarkably conserved and show significant parallels between species as distant as humans and zebrafish. Beyond coding regions exhibiting divergence in both species, non-coding regions and lncRNAs in human sperm and lincRNAs in zebrafish sperm were particularly frequent. lnc/linc RNAs exert widespread regulatory functions on downstream gene expression (62), suggesting substantial regulatory activity in genes determining sperm function. Differences in lncRNA expression have also been reported in sperm of men with compromised fertility (42) where they regulate genes involved in cytoplasmic transport, biosynthetic processes and spermatogenesis. Notably, we found little overlap with geno-informative markers previously identified in mammals, such as mouse, macaque and cattle (7).

Our genomic and proteomic data indicate that this gametic stage may play a key role in favouring high-functioning genomes, with potential to reduce transmission of heritable disease and to improve overall health and fitness. While the genomic data highlight clear divergence between sperm pools, reduced abundance of proteins related to neurodevelopmental processes and cancer in high-quality sperm suggests a direct link between phenotypic sperm quality and genetic quality. A link between sperm fitness and offspring fitness has been demonstrated in zebrafish (5,41), and we provide the genomic architecture underlying these differences. Neurodevelopmental disorders point to early developmental differences, which may also explain the observed 5-7% difference in survival during early zebrafish embryo development (5). In contrast, cancer is an ageing-related heritable disease, and the reduction of proteomic cancer biomarkers in high-quality sperm aligns with our previous observation of higher reproductive fitness and increased lifespan in zebrafish sired by longer-lived sperm (41). The signals among variants contributing to Polygenic Scores further support a link between overall genetic sperm quality and human health.

We would like to mention several limitations of this study. Donor numbers in the genomic and proteomic analyses are modest (n = 4–5 per assay), which limits generalisability across genetic backgrounds, although the most reproducibly diverging genes recurred in at least six of nine donors. Moreover, the polygenic and likely epistatic architecture of sperm quality means that no single locus or protein should be regarded as deterministic. While the link between sperm-level selection and offspring lifespan is direct in zebrafish, in humans it remains inferential, resting on the documented age-related disease relevance of the genes and proteins identified here. Several leading candidates are pleiotropic, with established roles both in neurodevelopment and in adult tumour suppression and senescence, and the relative contributions of these functions to selection during the gametic phase should be a focus of future research. Finally, our analyses are based on standard reference annotations and standard ageing-related gene databases remain incomplete.

Within-ejaculate sperm selection has historically been studied through the lens of fertility and reproductive medicine. The data presented here suggest a novel complementary view, namely that this same selection, by depleting alleles and proteins associated with age-related disease before fertilization, contributes to the heritable component of human healthspan. Identifying the molecular mechanisms of this pre-fertilization filter offers a route to understanding why, in zebrafish, the longevity of a sperm predicts the longevity of the offspring it sires, and provides a framework for asking the same question in humans.

## Materials and Methods

### Sample collection from human donors

For the assessing the effects of sperm selection on sperm phenotypic quality, we used the samples from 35 self-reported healthy donors collected at the University of East Anglia, UK. For the genomics analyses, we included ejaculates from nine self-reported healthy volunteers with sperm concentration >15 M/ml: nine samples used for genomics analyses were collected at the Rigshospitalet Copenhagen, Denmark. For TMT proteomics analyses we used four human sperm ejaculates: two from the University of East Anglia (UEA), Norwich, UK; two from Cape Town, South Africa (ZA)). For the biomarker validation of OFLM4 and CTNG, we recruited an additional five donors from the University of East Anglia (UEA), Norwich, UK. The samples from University of East Anglia and Cape Town were collected under ethics permit no. ETH2324-0997 and the samples collected at the Rigshospitalet Copenhagen were collected under ethics permit no. H-17012149.

### Sperm selection assays

All samples were collected near the laboratory and allowed to liquefy for 30 minutes at 37°C prior to the selection assay. We used two different assays to select for sperm within ejaculates:

### Methyl Cellulose Assay

A 10 cm-diameter watch glass was filled with 1% (w/v) methylcellulose (Sigma, Denmark) up to a 4cm radius and 10ml total volume. Then, 300µl of raw ejaculate was placed in the centre of the glass (the ejaculate was allowed to sink and filled a circle of 0.5cm in the centre of the glass). For each donor, we used five 300 µl aliquots allocated into five watch glasses per donor in parallel to ensure sufficient cells in the edge population for subsequent whole genome sequencing. The samples were incubated in the watch glasses for two hours at 37°C under 5% CO2 supplementation. 4ml aliquots were then collected from the outer circle of each watch glass at a radius of 3-3.5 cm, 2.5-3cm or 2-2.5cm (depending on the distance travelled by the sperm starting at the radius from the further circle with >10 cells under the microscope for each male) and placed in a 15ml tube. All five edge samples collected from the outer circle were pooled for each donor to ensure high enough cell numbers for sequencing. We also collected a sample (5 ml) from the centre of each watch glass at least 0.5 cm above the bottom of the glass to avoid contamination with dead sperm and somatic cells. To extract the sperm from the methylcellulose, 1µl of cellulase was added for every 1ml of collected sample and the sample was mixed thoroughly on a shaking table for two hours at 50°C. The samples were then centrifuged, the supernatant removed, and the sperm pellets were stored at -80° C for subsequent DNA extraction.

### Swim Up Assays

For the phenotypic analyses, we collected a raw (R T0), washed (W T0), and sub-samples from swim ups following incubation (T0, T4, T8, T24, T48) and for the genomic analyses we collected a raw sample (T0) and sub-samples from swim ups following incubation (T4, T24 for all five donors and T8 (n = 1 donor), T24 and T48 (n = 3 donors).

For the Swim Up, aliquots of 1ml raw ejaculate were placed in 15ml centrifuge tubes at a 1:1 ratio of human tubal fluid (HTF) medium containing (in mM): 72.8 NaCl, 4.69 KCl, 0.2 MgSO4, 0.37 KH2PO4, 2.04 CaCl2, 0.33 sodium pyruvate, 21.4 sodium lactate, 2.78 glucose, 21 HEPES, 25 NaHCO3, adjusted to pH 7.4 with NaOH (Fisher, UK) and incubated for 2, 4, 8, 12, 24 and 48 hours at 37°C with 5% CO2 in the air. Motile sperm, at each time point, were recovered by direct swim up as follows: after flicking each tube to homogenise the sample, 1ml was aliquoted and placed under 4ml of HTF medium in a new 15ml centrifuge tube at 37°C for 60 min in 5% CO2 as described above. The top layer of the swim up (3.5ml) was collected and centrifuged at 700xg for 10-20 min at room temperature (RT) prior to resuspension in 2ml HTF. After assessing sperm concentrations of samples using image cytometry, the cells were pelleted, the supernatant removed, and the pellet stored at -80°C until further use.

### Sample prep and selection assay

We used samples from 35 donors for sperm selection by Swim Up assay and subsequent phenotypic analyses. All raw samples were split into three aliquots: one third of the raw sample was used directly for a swim up and the remaining 2/3 were washed and split into two aliquots and incubated for 4h and 24h, respectively. In the washing step, the samples were washed twice in HTF lacking HAS. The sample was centrifuged at 500g for 10 minutes and after removal of the supernatant, 3ml of HFT+ was added and the sample was split again into two labelled tubes (T4 and T24). Following incubation, the samples were centrifuged at 500g for 10 minutes to supernatant and perform a direct swim up to isolate motile cells as described above.

The swim ups were performed in an Eppendorf tube as follows: 1 ml of human tubal fluid (HTF+) medium containing (in mM): 72.8 NaCl, 4.69 KCl, 0.2 MgSO4, 0.37 KH2PO4, 2.04 CaCl2, 0.33 sodium pyruvate, 21.4 sodium lactate, 2.78 glucose, 21 HEPES, 25 NaHCO3 and 3.5% human serum albumin (HSA) (Fisher, UK) adjusted to pH 7.4 with HCL was added to the semen/sperm pellet. During the swim up assay, cells were allowed to swim up for 1h at 37°C with 5% CO2 supplementation after which we collected 700µl of the sample from the top of the Eppendorf tube.

We then assessed phenotypic traits including sperm concentration, motility and velocity, sperm viability, sperm morphology, chromatin and DNA integrity (see Supplementary Methods for details on each of these).

### Whole genome sequencing

DNA was extracted using a standard phenol:chloroform:isoamyl alcohol protocol (see Supplementary Methods for details on DNA extraction) from the sperm pools and prepared using a PCR-free library preparation kit (KAPA HyperPrep kit; Roche) for a standard insert size (400-500 bp), (see Supplementary Methods for details). Sequencing was conducted by the Earlham Institute, UK, on an Illumina NovaSeq6000 to generate 150 bp paired-end reads.

### Bioinformatic analyses of genome data

All scripts and step-by-step instructions are available on: https://github.com/alicegodden/spermpool.

### Allele frequency estimations

Following quality control (see Supplementary Methods for details), variant detection of the alignments of reads from the T0 raw ejaculate followed the best practice hard-filtering workflow recommended by GATK. In brief, GATK 4.9.1 Haplotype Caller was used to call the variants and filter the heterozygous paternal sites. Single nucleotide polymorphisms (SNPs) were filtered using SelectVariant. Alignment of the reads from T0 raw ejaculate and selected sperm pools were used to generate base count profiles using pileup2pro2 (https://github.com/douglasgscofield/gameteUtils). SAMtools v1.9 option -q30, -Q30 and d100000 was used to exclude sites with low quality, allowing for sites with deep coverage.

For the likelihood ration tests, we only retained sites with a total coverage of > 31x or more than 2*total mean coverage were excluded (mean coverage was 206.69, 200.68, 176.31, 174.9, 261.17, 221.03, 271.5, 239.1023 and 211.443 for Donors 1 to 9 respectively). The heterozygous sites and base count profiles of the T0 raw and high performance sperm pools were used for likelihood ratio tests using the script hetPoolLikelihoods.pl available at https://github.com/douglasgscofield/gameteUtils following Lynch et al (2014) (44). The most significant allelic differences between the T0 raw and T4/24 sperm pools were identified by the largest likelihood ratios. Briefly, the major and minor alleles were assigned according to counts within the T0 raw sperm pools. To identify statistically significant loci between T0 raw and T4/T8/T24/T48 sperm pools, the threshold for genome-wide significance was set to > 99.9th quantile (critical values Donor1: 17.08, Donor 2: 11.91, Donor 3: 15.36, Donor 4:11.60, Donor 5: 11.37, Donor 6: 11.30, Donor 7: 11.67, Donor 8: 11.37, Donor 9:11.6).

### Gene function analyses and direction of selection

To identify genetic elements residing at statistically significant loci, identified loci were mapped onto annotated Homo sapiens reference genome GRCh38. To uncover the biological significance of the genetic differences resulting from the comparisons between T0 raw and selected fit sperm pools, the resulting gene overlaps were then used for gene enrichment and gene length analysis using ShinyGO v0.75 (45).

To evaluate whether variants under selection were enriched for predicted deleterious effects, variant consequences were annotated using Ensembl Variant Effect Predictor (VEP). Variants were classified according to VEP IMPACT categories (HIGH, MODERATE, LOW and MODIFIER) and predicted molecular consequences. Putatively deleterious variants were defined as those with HIGH-impact consequences, including stop-gained, frameshift, splice donor/acceptor and start/stop-loss variants. For these variants, the alternative (ALT) allele was considered the deleterious allele. Differences in ALT allele frequency (ΔALT) between groups were compared between deleterious and non-deleterious variants (MODIFIER and synonymous categories) using Wilcoxon rank-sum tests to assess whether putatively deleterious alleles were preferentially depleted. Owing to the predominance of non-coding variants among the divergent loci, this categorical analysis was interpreted as a complementary assessment of deleteriousness.

We also assessed polygenic scores for variants in our dataset. VCF files were generated from each sperm pool BAM file with a bcftools v1.15.1 (46) pipeline as follows: bcftools mpileup -a AD,ADF,ADR -r “${ncbi_chr}” -q 30 -Q 30 -f GRCh38_latest_genomic.fna “${bam_file}” | bcftools call -mv -f GQ -Oz -o “$output_file”. Chromosome files were concatenated, and chromosome identifiers converted to Ensembl format. To allow integration with the Pan-UK Biobank dataset, the VCF files were lifted over to the GRCh37 reference genome coordinate system using GATK v4.6.0.0, with Picard LiftoverVCF. The GRCh37 reference genome was obtained from Ensembl, Homo_sapiens.GRCh37.dna_sm.primary_assembly.fa.gz, version 27/11/2015. The chain file GRCh37_to_GRCh38.chain.gz, version 25/07/2014, https://ftp.ensembl.org/pub/assembly_mapping/homo_sapiens/ was downloaded from Ensembl. Polygenic Score (PGS) data were obtained from the Pan-UK Biobank phenotype manifest: https://docs.google.com/spreadsheets/d/1AeeADtT0U1AukliiNyiVzVRdLYPkTbruQSk38DeutU8/edit?gid=1450719288#gid=1450719288 in May 2025. The data were filtered to retain CHR, POS, REF, ALT and beta_EUR columns, with the latter representing the effect size of the alternative allele as estimated in UK Biobank participants of inferred European ancestry. We extracted information on allele frequency, chrom, pos, ref and alt columns from the sperm pool VCF files and retained all positions that overlapped with variants in the Pan-UK Biobank dataset. The sperm pool allele frequencies and beta_EUR of all overlapping loci for a phenotype were multiplied and the product for each phenotype was added up, to obtain a score for each donor.

### Comparison between human and zebrafish data

To identify conserved genetic pathways between zebrafish and human sperm pools selected for fitness, we reanalysed the zebrafish sperm pool data from Alavioon et al. (2017) (5). To process the genomic data, we used the latest zebrafish GRCz11 Danio rerio reference genome to align the genomic reads and employed the same analysis pipeline and hard filtering as for human samples (see Supplementary Methods for details). To further minimise potential sequencing bias in the zebrafish data, the cut-off point for genome-wide identification of significant loci in the zebrafish data was set to the 99.9th quantile; critical values were 13.99, 26.51, 10.39 for male ID: Male_31, Male_32 and Male_34, respectively. We then used Venn diagrams on significant genes by converting zebrafish genes into human gene homologs to identify genes shared between humans and zebrafish and performed GO analyses using ShinyGO v0.75 (45) on the 153 shared genes.

### Sperm selection assays for TMT proteomics

Samples from four donors in total, two for each assay were used. The samples were washed in human tubal fluid (HTF) and resuspended in 200 µl HTF. We collected two samples representing a less fit (Centre/T0) and a fitter (Edge/T4) sperm pool from each donor from Methyl Cellulose and Swim Up (assays as described above; see Supplementary Methods for exact details) and stored them at -80°C.

### Protein extraction and proteomics

For protein extraction, samples were thawed, and 450 ml of acetone was added to each of the thawed samples. Samples were vortexed for 15 minutes, then centrifuged at 10,000 rpm for 10 minutes. Supernatant was then removed, and 500 ml of acetone added to each sample. Each sample was then spun gently for two minutes manually then centrifuged for 15 mins at 10000 rpm. Supernatant was removed and samples were air dried for 1 and a half hours.

Protein pellets were resuspended in 100 µl of 2.5% sodium deoxycholate (SDC; Merck) in 0.2 M EPPS-buffer (Merck), pH 8.5, and the samples vortexed under heating. The samples were treated with dithiothreitol and iodoacetamide to alkylate cysteine residues and digested with trypsin in the SDC buffer according to standard procedures. After the digest, the SDC was precipitated by adjusting to 0.2% TFA, and the clear supernatant subjected to C18 SPE (Reprosil-Pur 120 C18-AQ, 5 um, Dr. Maisch GmbH, Germany). Peptide concentration was estimated by running an aliquot of the digests on LCMS (see below). TMT labelling was performed using 8 channels from a TMT™10plex kit (TMT lot: VH306773, ThermoFisher Scientific, Hemel Hempstead, UK) according to the manufacturer’s instructions with slight modifications; samples were dissolved in 90 µl of 0.2 M EPPS buffer (MERCK)/10% acetonitrile, and 200 µg TMT reagent dissolved in 22 µl of acetonitrile was added.

Samples were assigned to the TMT channels in two separate experiments, according to the table below. After 2 h incubation, aliquots of 1.5 µl from each sample from each individual TMT experiment were combined in 250 µl 0.2% TFA, desalted, and analysed on the mass spectrometer to check the labelling efficiency and estimate total sample abundances. The main sample aliquots were quenched by adding 8 µl of 5% hydroxylamine and combined to roughly level abundances. The peptides were desalted using a C18 Sep-Pak cartridge (200 mg, Waters, Wilmslow, UK). The eluted peptides were dissolved in 500 µl of 25 mM NH4HCO3 and fractionated by high pH reversed phase HPLC. The samples were loaded to an XBridge® 3.5 µm C18 column (150 x 3.0 mm, Waters). Fractionation was performed on an ACQUITY Arc Bio System (Waters) with the following gradient of solvents A (water), B (acetonitrile), and C (25 mM NH4HCO3 in water) at a flow rate of 0.5 ml min-1: solvent C was kept at 10% throughout the gradient; solvent B: 0-5 min: 5%, 5-10 min: 5-10%, 10-60 min: 10-40%, 60-75 min: 40-80%, followed by 5 min at 80% B and re-equilibration to 5% for 24 min. Fractions were collected every 1 min and concatenated by combining fractions of similar peptide concentration to produce 20 final fractions for MS analysis. A replicate of one donor sample (two TMT channels) was fractionated using the Pierce™ High pH Reversed-Phase Peptide Fractionation Kit (Thermo) to produce seven fractions.

### TMT analyses

Aliquots of all fractions were analysed by nanoLC-MS/MS on an Orbitrap Eclipse™ Tribrid™ mass spectrometer equipped with a FAIMS Pro Duo interphase coupled to an UltiMate® 3000 RSLCnano LC system (Thermo Fisher Scientific, Hemel Hempstead, UK). The samples were loaded onto a trap cartridge (PepMap™ Neo 5 µm C18 300 µm X 5 mm Trap Cartridge, Thermo) with 0.1% TFA at 15 µl min-1 for 3 min. The trap column was then switched in-line with the analytical column (Aurora Frontier TS, 60 cm nanoflow UHPLC column, ID 75 µm, reversed phase C18, 1.7 µm, 120 Å; IonOpticks, Fitzroy, Australia) for separation using the following gradient of solvents A (water, 0.1% formic acid) and B (80% acetonitrile, 0.1% formic acid) at a flow rate of 0.23 µl min-1 : 0-3 min 1% B (parallel to trapping); 3-10 min linear increase B to 9 %; 10-105 min increase B to 50%; followed by a ramp to 99% B and re-equilibration to 1% B.

Data were acquired with the following parameters in positive ion mode: MS1/OT: resolution 120K, profile mode, mass range m/z 400-1600, AGC target 4e5, max inject time 50 ms, FAIMS device set to three compensation voltages (-35V, -50V, - 65V) for 1 s each; MS2/IT: for each CV, data dependent analysis with the following parameters: 1 s cycle time Rapid mode, centroid mode, quadrupole isolation window 0.7 Da, charge states 2-5, threshold 1.9e4, CID CE = 33, AGC target 1e4, max. inject time 50 ms, dynamic exclusion 1 count for 15s with mass tolerance of 7 ppm; MS3 synchronous precursor selection (SPS): 10 SPS precursors, MS2 isolation window 0.7 Da, HCD fragmentation with CE=65, Orbitrap Turbo TMT and TMTpro resolution 30k, AGC target 200%, max inject time 100 ms; Real Time Search (RTS): protein database UP000005640_human_2024_9606.fasta (Uniprot, 20,609 entries), enzyme trypsin, 1 missed cleavage, oxidation (M) as variable, carbamidomethyl (C) and TMT6plex as fixed modifications, precursor tolerance 10 ppm, Xcorr = 1.4, dCn = 0.1.

### Proteome data analyses

Scripts are on: https://github.com/alicegodden/spermpool.

The acquired raw data were processed and quantified in Proteome Discoverer 3.1 (Thermo Fisher Scientific); all mentioned tools of the following workflow are nodes of the proprietary Proteome Discoverer (PD) software. The UP000005640_human_2024_9606.fasta protein fasta database (Uniprot, 20,609 entries) was imported into PD adding a reversed sequence database for decoy searches. The database search was performed using the incorporated search engines CHIMERYS (MSAID, Munich, Germany) and Comet. The processing workflow for both search engines included recalibration of MS1 spectra (RC), reporter ion quantification by most confident centroid (20 ppm) and a search on the human protein database (as imported above). For CHIMERYS the Top N Peak Filter was applied with 20 peaks per 100 Da. Then the inferys_3.0.0. fragmentation prediction model was used with fragment tolerance of 0.3 Da, enzyme trypsin with 1 missed cleavage, variable modification oxidation (M), fixed modifications carbamidomethyl (C) and TMT6plex on N-terminus and K. For Comet the version 2019.01 rev.0 parameter file was used with default settings except precursor tolerance set to 6 ppm and trypsin missed cleavages set to 1. Modifications were the same as for CHIMERYS.

The consensus workflow included the following parameters: intensity-based abundance, normalisation on total peptide abundances, protein abundance-based ratio calculation, only unique peptides (protein groups) for quantification, TMT channel correction values applied (Lot VH306773), co-isolation/SPS matches/CHIMERYS Coefficient thresholds 50%/70%, 0.8, missing values imputation by low abundance resampling, hypothesis testing by t-test (background based), adjusted p-value calculation by BH-method. The results were exported into a Microsoft Excel table including data for normalised and un-normalised abundances, ratios for the specified conditions, the corresponding p-values and adjusted p-values, number of unique peptides, q-values, PEP-values, identification scores from both search engines; FDR confidence filtered for high confidence (strict FDR 0.01) only.

We used AlphaFold (47–49) to generate molecule views of the candidate proteins.

### Validation of OLFM4 and CTNG

Following sample collection, liquefaction and CASA analysis to assess sperm motility in each sample, samples were aliquoted to approximately 1.5 million cells in 1 mL of human tubal fluid (HTF) medium. Aliquots were incubated for 4 hours at 37 °C with 5% CO2 prior to antibody staining. Following incubation, sperm samples were reassessed for motility and cell number using CASA, then immediately stained with eFluor™ 520 fixable viability dye (eBioscience, Invitrogen) according to the manufacturer’s protocol. After viability staining, cells were fixed in 4% paraformaldehyde (PFA) at 4 °C.

Fixed cells were then blocked using Human Fc Receptor Blocking Solution (eBioscience, Invitrogen) to minimise non-specific antibody binding. Fluorophore-conjugated primary antibodies were subsequently added and incubated under standard conditions. The following antibodies were used: anti-OLFM4 clone 12, conjugated to R-phycoerythrin (PE), at a 1:10 dilution (Novus Biologicals, NBP3-06415PE); anti-CTNG (CTNG/2155R), conjugated to mFluor Violet 450 SE, at 1:10 dilution (Novus Biologicals, NBP3-08485MFV450); and anti-AOC3 (VAP1/AOC3 - TK8-14), conjugated to Alexa Fluor 594, at 1:10 dilution (Bio-Techne, NBP2-81043AF594.). Throughout staining, cells underwent minimal washing steps to preserve cell numbers and were stored overnight at 4 °C prior to flow cytometry analysis.

Samples were analysed using the BD FACSDiscover™ S8 Cell Sorter (BD Biosciences), operated in fully spectral mode. No cell sorting was performed. Imaging and fluorescence data were acquired simultaneously for each sample with fixable viability dye (eFluor™ 520) detected using imaging laser B1 to allow for imaging. Events were gated to isolate the sperm population (P1), based on forward and side scatter properties. All subsequent data acquisition and analysis were restricted to the P1 population. For each antibody-stained sample, between 5,000 and 50,000 individual events within the P1 gate were recorded, depending on sample quality and cell availability. Spectral unmixing was performed using BD FACSChorus™ software (BD Biosciences), based on single-stained controls for each fluorophore-conjugated antibody and the fixable viability dye. An unstained sperm sample was included in experiment set to control for cellular autofluorescence during unmixing. Compensation and unmixing matrices were applied prior to further analysis.

Flow cytometry image files were merged with the primary fluorescence data using BD CellView™ Image Extractor. The resulting datasets were processed in R v4.4.3 (50). Data were normalised using the CytoNorm package (51), and quality control was performed using PeacoQC (52) to identify and remove anomalous events. Gated and cleaned data were subsequently analysed in FlowJo v 10.10.0 (BD Life Sciences - FlowJo, LLC, Ashland, OR, USA. https://www.flowjo.com), with all events restricted to the P1 sperm population (see Suppl. Fig. S5-2 for gating strategy).

Statistical modelling of marker expression in live and dead cell populations was conducted in R v4.4.3 (50), with residual diagnostics simulated using the DHARMa package (53) to evaluate model fit and distribution assumptions. Model summaries were formatted using the Stargazer package for publication (54).

### Data availability

All raw datasets have been uploaded to respective platforms. The sperm phenotypic data, R scripts associated with phenotype statistical analyses, the list of significantly differentially expressed genes form the human genomic analyses and the list of significantly differentially expressed proteins can be found on dryad (doi:10.5061/dryad.4qrfj6qkn). The data files containing the raw whole-genome sequencing data for all human sperm pools can be found in the European Nucleotide Archive under accession number PRJEB80525. The mass spectrometry proteomics data have been deposited to the ProteomeXchange Consortium via the PRIDE partner repository with the dataset identifier PXD056057 and 10.6019/PXD056057; PXD056228 and 10.6019/PXD056228; and PXD056234 and 10.6019/PXD056234””.

## Acknowledgments

We thank Carlo Martins for his help with the proteomics work and Sally Otto and Cock van Oosterhout for feedback on earlier drafts of this manuscript. This research was supported by grants from the European Research Council (ERC, HapSelA-336633 and SELECTHAPLOID-101001341) and Natural and Environmental Research Council (NERC, NE/S011188/1) to SI and a BBSRC PhD studentship to DM.

